# When More Is Not Better: Increased Motor Cortex Recruitment In Older Adults Is Associated With Performance Decline During High-Demand Cognitive-Motor Tasks

**DOI:** 10.64898/2026.04.24.720565

**Authors:** Marine Keime, Sanskar Ranglani, Monika Harvey, Cassandra Sampaio-Baptista

## Abstract

Age-related changes in brain structure and function lead to reorganisation in neural activity during cognitive-motor tasks. A central question is whether increased brain activity in older adults reflects compensatory mechanisms that help maintain performance, or inefficient recruitment due to neural dedifferentiation. In this context, the Compensation-Related Utilization of Neural Circuits Hypothesis (CRUNCH) proposes that older adults recruit more brain resources as task demands rise, but reach a limit beyond which this no longer supports performance. This study examined brain activity changes in older adults in response to increased cognitive-motor task demands.

Twenty-one right-handed older adults performed an Action Selection (AS) task with three difficulty levels during functional MRI. Behavioural performance declined with increasing task difficulty. Whole-brain analyses revealed that higher task complexity was associated with increased activation in sensorimotor and prefrontal regions, particularly the left dorsolateral prefrontal cortex (DLPFC) and left premotor cortex (PMd). Importantly, higher activity in subcortical regions (caudate nucleus) was associated with better performance at lower difficulties, while greater activation in cortical motor areas (primary motor cortex (M1), and the supplementary motor area (SMA)) was linked to worse performance at high difficulty. Additionally, greater increases in sensorimotor activity (M1, PMd, DLPFC) across difficulties correlated with greater declines in performance, supporting the notion of inefficient resource recruitment. Together, the findings support a novel hypothesis, compatible with CRUNCH, of a subcortical to cortical shift (SCOS) with increasing cognitive-motor demand, where the transition from efficient subcortical processing to over-reliance on cortical motor areas may contribute to performance decline in ageing.

## 1. INTRODUCTION

Ageing is accompanied by structural (Fujiyama et al., 2016), functional (Wu et al., 2007) and metabolic (Seidler et al., 2010) changes in the brain, which collectively lead to widespread reorganisation of neural networks. Compared to younger adults, older adults often exhibit increased and widespread brain activity during task execution, particularly during cognitive and motor tasks. For example, the primary motor cortex (M1) appears to play a diminished role in motor control with age, while regions such as the premotor cortex (PMd) and dorsolateral prefrontal cortex (DLPFC) show increased recruitment (Goble et al., 2010; Heuninckx et al., 2008, 2005; Santos Monteiro et al., 2017; Tscherpel et al., 2020; Verstraelen et al., 2020). However, it is still debated whether this increased activity represents compensatory reorganisation that supports performance, or instead dedifferentiation, reflecting reduced neural specificity and efficiency (Davis et al., 2008; Maes et al., 2020; Rurak et al., 2021; Tak et al., 2021).

Two prominent theories have attempted to explain age-related changes in brain activity. The Posterior-Anterior Shift in Ageing (PASA) proposes that frontal regions are recruited to compensate for declining function in posterior areas, such as the occipital and parietal cortices (Davis et al., 2008; Michely et al., 2018; Tscherpel et al., 2020). As such, some studies have reported a positive link between increased DLPFC activity and task performance in older adults (Davis et al., 2008; Solé-Padullés et al., 2006). In parallel, the Hemispheric Asymmetry Reduction in Older Adults (HAROLD) model suggests that older adults increasingly engage both hemispheres to maintain performance on tasks that typically show lateralized activity in younger adults (Cabeza, 2002). Both theories were originally developed in the context of cognitive processes such as memory and have been less thoroughly tested in cognitive-motor domains (Heuninckx et al., 2005; Hutchinson et al., 2002; Mattay et al., 2002; Michely et al., 2018; Riecker et al., 2006).

Crucially, increased brain activity is not always associated with better performance. Recent evidence indicates that elevated activity in core motor areas (M1, SMA, PMd) may not be beneficial and could instead reflect network inefficiency or degradation (Knights et al., 2021; Morcom and Henson, 2020; Tscherpel et al., 2020). Knights and colleagues (2021), for instance, demonstrated through a multivariate Bayesian analysis that increased ipsilateral motor activity carried no additional information about the motor task compared to the contralateral hemisphere. These alterations may instead reflect reduced specificity of neural responses, leading to the nonselective engagement of broader brain regions (Li et al., 2015; Morcom and Henson, 2020; Tscherpel et al., 2020).

To reconcile these conflicting findings, the Compensation-Related Utilization of Neural Circuits Hypothesis (CRUNCH) was proposed (Reuter-Lorenz & Cappell, 2008). CRUNCH suggests that older adults initially recruit additional neural resources under low-to-moderate task demands to maintain performance, but eventually reach a ceiling beyond which further recruitment is ineffective. A recent review on motor control and ageing supports the idea that increased activation in prefrontal and motor regions is modulated by task complexity (Zapparoli et al., 2022). However, empirical evidence supporting CRUNCH in motor domains remains limited (Van Ruitenbeek et al., 2023), and it is also unclear whether such activity reflects true compensation or inefficient processing. In the present study, we investigate this latter aspect, focusing on within-subject modulation in older adults, given that prior work has already characterised age differences (Zapparoli et al., 2022; Van Ruitenbeek et al., 2023). Instead, older adults act as their own controls across task demands, thus allowing compensatory capacity to be evaluated through changes in neural activity with increasing difficulty, rather than directly testing age-related differences relative to younger populations.

More specifically, the present study aimed to investigate whether demand-related changes in brain activity during a cognitive-motor task reflect compensation or inefficiency in older adults. Using a within-subject fMRI design, brain activity and behavioural performance were examined during an Action Selection (AS) task with three levels of difficulty. Based on the CRUNCH, the study addressed the following hypotheses:

### Hypothesis 1

activity in predicted regions (SMA, M1, PMd, DLPFC) will increase with difficulty.

### Hypothesis 2

higher activity in SMA, M1 and PMd will be linked to poorer performance in higher difficulty levels, reflecting inefficient recruitment, whereas higher activity in DLPFC will be associated with better performance, as a sign of compensatory recruitment.

### Hypothesis 3

greater activity increases in SMA, M1, PMd, and DLPFC across difficulty levels will be linked to lower performance decline across difficulty levels, potentially reflecting compensatory neural adjustment.

## 2. METHODS

### 2.1 Participants and design

21 older adults (> 55 years old; mean age = 68.76, sd = 8.72, 13F), right-handed (assessed by the Edinburgh Handedness (Oldfield, 1971)), and MRI safe, were recruited for this study. Participants were screened for cognitive decline using the Montreal Cognitive Assessment (scoring > 26, Nasreddine et al., 2005). Data was collected at the Centre of Cognitive Neuroimaging, University of Glasgow. The study was approved by the College of Medical Veterinary and Life Sciences Ethics Committee (ref: 200220405).

This study employed a within-subject design. Participants came in for two sessions: one for behavioural training on the AS task and one MRI session during which they executed the task inside the scanner as well as prior to the scan, as a refresher (Fig. 1). The order of difficulty during the training and the refresher was randomised. During the MRI session, task difficulty was presented in a fixed increasing order (1, 2, 3) to ensure a consistent progression of cognitive demand across participants and avoid order effects.

**Figure 1:**
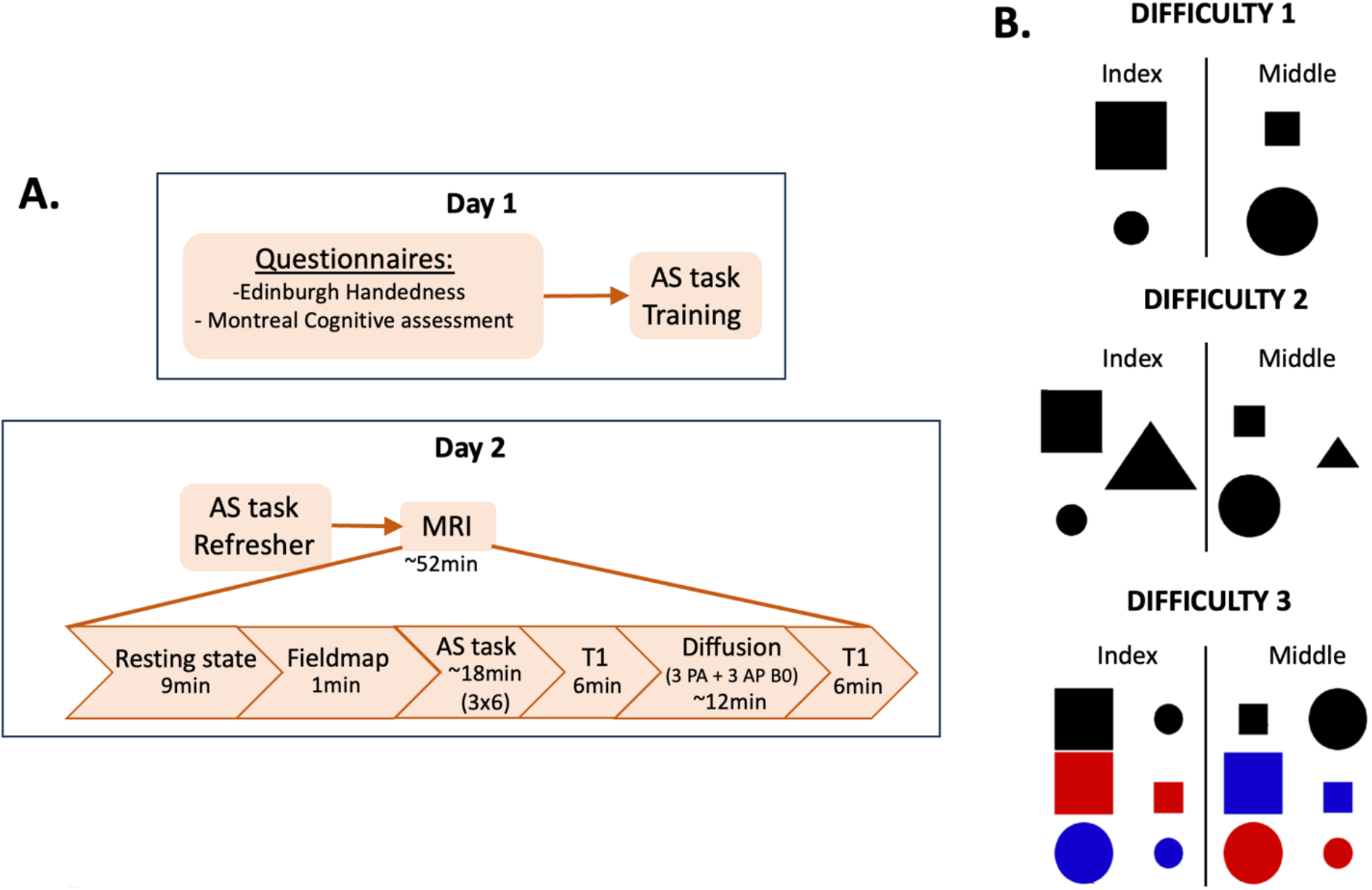
Study design and procedure. A. Day 1 consisted in training participants on the AS task until performance plateaued (RT slope < 0.15), as well as fill out some questionnaires. Day 2 consisted of executing the AS task inside the MRI scanner. Participants got a refresher of the AS task before going into the scanner also. B. Illustration of the AS block with increasing difficulties. When a shape presented on the left side of the instruction screen appeared, the participant had to press a button with their index finger, whereas if a shape from the right side of the screen appeared, they had to press a button with their middle finger. Each difficulty consisted of two blocks of EO, AS, and VO (inside the scanner) each including 25 trials (6 blocks total, repeated for 3 difficulties). Shapes were presented for 1sec, separated by a fixation cross (∼500ms). Task ran on Matlab (2020a), using Psychtoolbox packages.

### 2.2 Action Selection task

The behavioural outcome of this study was measured by the AS task (Johansen-Berg et al., 2002; O’Shea et al., 2007; Rushworth et al., 2003). This task was modified, and two higher difficulty levels were developed. The new difficulty levels were piloted on six older adults and selected from a set of four options, based on statistical and observational differences between difficulties (see Supplementary materials 1).

The task required participants to press a button as quickly as possible in response to shapes presented at the centre of the screen. It included three block types: Execution Only (EO), View Only (VO), and Action Selection (AS). In EO blocks, participants pressed the same key on every trial: the middle finger in the first block and the index finger in the second, regardless of shape. VO blocks, presented only in the scanner, required participants to observe the shapes without responding. In AS blocks, participants pressed either the index or middle finger according to the shape, with three escalating levels of difficulty.

Difficulty 1 required pressing the index finger for a big square or small circle, and the middle finger for a big circle or small square. Difficulty 2 included the same rules as difficulty 1, with the addition of a triangle cue: participants pressed the index finger for a large triangle and the middle finger for a small triangle. Difficulty 3 incorporated colour: black shapes followed the difficulty 1 rules, while colourful shapes required the index finger for red squares or blue circles, and the middle finger for red circles or blue squares (all sizes; Fig. 1).

Each difficulty included two blocks of VO, EO, and AS, presented in a fixed order, with 25 trials per block. Shapes were displayed for 1 s, separated by a 500 ms fixation cross, giving 1.5 s per trial. Participants were trained on the task the day before scanning until their response times plateaued (RT slope < 0.15) to minimize learning effects. All responses were made using a right-hand button box.

### 2.3 Behavioural analysis

AS performance was measured for each participant as the RT cost (sec), calculated by subtracting the mean RT in EO blocks from the mean RT in AS blocks. Incorrect and missed trials were excluded from the RT cost calculation.

Additionally, as this study involved a task at high complexity, it was crucial to take errors into consideration for the measure of AS performance. Hence, the inverse efficiency score (IES) was also used as a performance measure, dividing the RT cost by the proportion of correct trials (Bruyer and Brysbaert, 2011). All results using the IES as a performance measure can be found in the Supplementary materials 2. Overall, the IES results showed a similar pattern to the RT costs, with performance declining at higher difficulty levels and greater cortical motor activation associated with poorer performance at high difficulty.

A one-way repeated measures ANOVA was conducted to test for the main effect of difficulty, followed by Bonferroni corrected post-hoc two-tailed paired t-tests between each difficulty.

### 2.4 MRI acquisition

Imaging was performed on a 3.0T Siemens Magnetom MRI system (Siemens, Erlangen, Germany). A task-fMRI was acquired with the following sequence parameters: Multiband sequence, acceleration factor 4, 80 slices, 2×2×2 mm^3^ voxel size, TR = 1200 ms; echo time = 26.0 ms; flip angle = 60°, mmview = 192×192mm. Two whole-brain T1 echo planar images were acquired for registration purposes (192 slices, 0.8×0.8×0.8 mm^3^ voxel size, TR = 2300 ms; echo time = 3.15 ms; flip angle = 9°, field of view = 255×255), in addition to a field map (46 slices, 3.7×3.7×3.7 mm^3^ in plane resolution, TR = 488 ms, difference in echo time = 2.46, flip angle = 60°, field of view = 256×256mm).

### 2.5 MRI data analysis

One participant was excluded from analyses involving difficulty 3 due to them not responding to any trial. Data from this participant was not excluded for analyses involving difficulty 1 and/or 2 only.

#### 2.5.1 FMRI pre-processing

FSL was used to analyse all fMRI data. FEAT was used to pre-process and analyse the fMRI data (Woolrich et al., 2004, 2001). The first EPI volume was deleted. The high-resolution image was used as an expanded functional registration, in addition to the structural (FLIRT, Jenkinson et al., 2002) and standard registrations (FNIRT, (Andersson, Jenkison & Smith, 2007). Nonbrain structures were removed using BET (Smith, 2002). Motion was corrected using MCFLIRT (Jenkinson et al., 2002).

### 2.5.2 FMRI first-level analysis

First-level designs were fitted to each AS difficulty runs using FEAT (Woolrich et al., 2001). A double gamma-HRF convolved boxcar regressor and its temporal derivative was used. The View Only (VO) blocks were used as baseline, the statistical design included Execute Only (EO) and AS contrasts, as well as AS > EO. Standard and extended motion parameters were used, adding an extra 18 regressors of no interest, to correct for head motion. Five participants mistakenly performed the AS instead of the EO task in one block of one difficulty. Those blocks were hence cut out from the data, and the statistical design was readapted to the new timings.

### 2.5.3 Whole-brain voxel-wise FMRI higher-level analyses

To test hypothesis 1, main effects of difficulty level were investigated (1 vs 2, 1 vs 3, 2 vs 3). In addition, a map of the average whole-brain BOLD activity (AS > EO) for each difficulty was calculated to assess general AS effects. The contrasts 1 and -1 were applied to extract both the positive and the negative mean activity during AS vs EO. Results of the average brain activity during each AS task difficulty can be found in Supplementary materials 2.

To test hypothesis 2, a voxel-wise whole-brain correlation analysis was conducted within a general linear model, relating AS > EO BOLD activity to AS performance (demeaned RT cost), separately for each difficulty. Demeaned performance was entered as a regressor, and 1 and −1 contrasts were used to test for positive and negative relationships, with demeaned age included as a covariate of no interest. The analysis was also carried out using demeaned IES as a performance measure (see Supplementary materials 2).

To test for hypothesis 3, AS > EO activity of the three task scans (for each participant) was modelled in a second level analysis weighing 1, 2, and 3 for each difficulty. Using fixed effects, 1 and -1 contrasts were applied to get both the linear increase and the linear decrease in activity. The linear increase was then input into a higher-level analysis, regressing for age and correlating with the RT cost linear slope (demeaned). Similarly, 1 and -1 contrasts were applied to get positive and negative correlations between activity change and RT cost change. The analysis was also carried out using demeaned IES slope, which can be found in supplementary materials 2.

Random effects were applied using FLAME to the higher-level analyses. Group Z (Gaussianized T) statistic images were thresholded using clusters determined by Z > 2.3 and a corrected cluster extent significance threshold of p < 0.05.

#### 2.5.4 Region of interest analysis

For completeness, the above analyses were also tested at a regional level. Based on whole-brain results, we specifically tested for effect in left M1 (LM1), left PMd (LPMd), SMA and left DLPFC (LDLPFC). Details regarding the analyses done and the results can be found in Supplementary materials 2.

## 3. RESULTS

### 3.1 AS performance declines with increasing difficulty

We tested for differences in RT cost between difficulties, and as expected, the main effect of difficulty was significant (repeated-measures ANOVA F(2,38) = 26.07, p < 0.0001, Fig. 2.A). Bonferroni post-hoc tests revealed RT cost was significantly higher in difficulty 3 vs 1 (paired t-test, t(19) = -5.95, pcorr < 0.0001, Fig. 2.A), as well as in difficulty 3 vs 2 (paired t-test, t(19) = -5.45, pcorr < 0.0001, Fig. 2.A). No difference was found between difficulty 1 and 2 (paired t-test, t(19) = -0.613, pcorr = 0.57).

**Figure 2:**
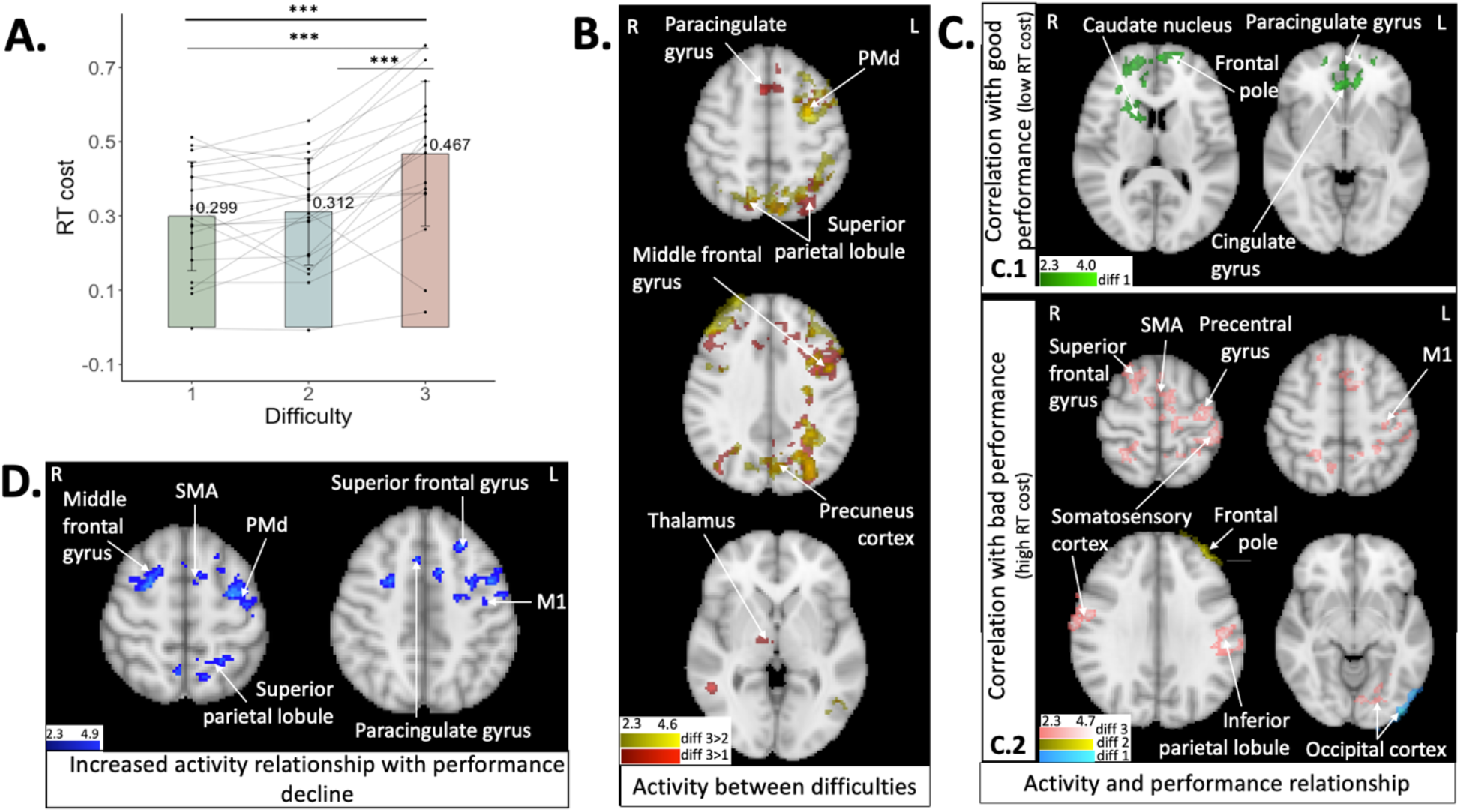
Task performance and brain activity. A. RT cost (sec) between difficulties. Main effect of difficulty (repeated-measures ANOVA, F(2,38) = 26.07), p < 0.0001, N = 20). Individual bar charts represent means, error bars are standard deviations and dots are individual data points. Full thick lines illustrate significant main effects of difficulty, full thin lines represent significant post-hoc comparisons. Significant p-value of ≤ 0.001 represented by ***. B. Hypothesis 1. BOLD activity difference between difficulties (all p < 0.05 corrected, N = 20). Red gradient illustrates activity higher in difficulty 3 vs 1, yellow is activity higher in 3 vs 2. C. Hypothesis 2. Correlation analysis between whole-brain activity (AS > EO) and AS performance (RT cost) (all p < 0.05 corrected, N = 21 for difficulty 1 & 2, N = 20 for difficulty 3). B.1. BOLD activity correlated with better performance (low RT cost) for difficulty 1 (green gradient). B.2 BOLD activity correlated with poorer performance (high RT cost) (cyan gradient for difficulty 1, yellow for difficulty 2, pink for difficulty 3). D. Hypothesis 3. Correlation analysis between whole-brain activity increase slope (AS > EO) and the AS performance slope (RT cost linear slope over difficulties). Blue gradient represents a positive correlation between activity slope and RT cost slope (higher activity increase linked to higher RT cost increase). N = 20, all p < 0.05 corrected.

### 3.2 Activity in sensorimotor and frontal networks during the AS task increases with higher difficulty

To address the first hypothesis, we tested for differences in whole-brain activity between difficulties. We expected increases in SMA, M1, PMd and DLPFC with increasing difficulty. No differences were found between difficulty 2 and 1. Activity was greater in difficulty 3 compared to both difficulty 1 and 2 in LPMd, bilateral superior parietal lobule, precuneus and occipital cortex (all p < 0.05, corrected; Fig. 2.B). Activity in the left middle and frontal gyrus (DLPFC) was also significantly greater in difficulty 3 vs 1 and 2, and was spatially wider in the 3 vs 1 contrast. The left supramarginal gyrus showed greater activation in difficulty 3 vs 2, while the paracingulate gyrus and right thalamus were more activated in difficulty 3 vs 1. No brain regions showed greater activation in the lower difficulty conditions compared to higher difficulty (1 > 2, 1 > 3, 2 > 3).

Therefore, higher activity in sensorimotor and frontal network (LPMd and LDLPFC specifically) were found with the highest difficulty (but not SMA and M1), partially aligning with the main hypothesis.

### 3.3 Motor planning areas linked to better AS performance at low difficulty and poorer performance at high difficulty

When examining whole-brain correlations with AS performance (RT cost) for hypothesis 2, we expected greater activity in SMA, M1, and PMd to be associated with poorer performance in higher difficulty levels (Knights et al., 2021; Morcom and Henson, 2020; Tscherpel et al., 2020), whereas higher DLPFC activity would be associated with better performance. Activity in the frontal pole, right caudate nucleus (subpart of the basal ganglia), paracingulate gyrus and cingulate gyrus were found to be correlated with better performance in difficulty 1 (all p < 0.05 corrected, Fig. 2.C.1). No brain areas were found to be correlated with better performance in difficulty 2 and 3.

On the other hand, poorer performance in the AS task (high RT cost) was correlated with activity in the occipital cortex in difficulty 1 (all p < 0.05 corrected, Fig. 2.C.2). In difficulty 2, activity in the left frontal pole was associated with poorer performance (all p < 0.05 corrected, Fig. 2.C.2). Lastly, poorer performance was associated with activity in the right superior frontal gyrus (DLPFC), bilateral parietal lobule, SMA, left somatosensory cortex, precentral gyrus, M1 and occipital cortex in difficulty 3 (all p < 0.05 corrected, Fig. 2.C.2).

Overall, higher activity in motor planning areas in the basal ganglia were associated with better performance at low difficulty levels, whereas cortical motor regions (SMA, LPMd, LM1) were associated with poorer performance at high difficulty, consistent with the prediction. However, contrary to the hypothesis, higher DLPFC activity was also linked to poorer performance, providing no evidence for a compensatory prefrontal effect.

### 3.4 Increases in motor regions activity with difficulty linked to greater performance decline

For the final hypothesis, we expected greater increases in activity in M1, LPMd, SMA, and DLPFC across difficulty levels to be associated with reduced performance decline. We tested for correlations between changes in brain activity with changes in RT cost between difficulties.

Instead, increases in LPMd, SMA, LM1, bilateral middle and superior frontal gyri (DLPFC), left superior parietal lobule, and the paracingulate gyrus were positively correlated with increases in RT cost slope (all p < 0.05 corrected, Fig. 2.D). Therefore, greater activity change in these regions across difficulties correlated with greater declines in performance. No negative correlations were observed, indicating that activity increases were not linked to lower performance decline across the difficulty levels.

## 4. DISCUSSION

This study aimed to investigate whether task demand-related changes in brain activity reflect compensation or inefficiency in older adults. AS performance declined with increasing task difficulty (Fig. 2.A, Supplementary Fig. S3). Successful performance under low-demand conditions was supported by subcortical activation, whereas greater difficulty elicited increased cortical motor (as well as parietal and prefrontal) involvement that was also associated with worse performance.

In line with predictions, activity in DLPFC and PMd increased with difficulty, as shown in both whole-brain and region of interest analyses (Fig. 2.B, Supplementary Fig. S5). This is consistent with previous findings showing greater recruitment of frontal and premotor areas in older adults during motor and cognitive-motor tasks (Goble et al., 2010; Heuninckx et al., 2008, 2005; Santos Monteiro et al., 2020; Tscherpel et al., 2020; Verstraelen et al., 2020). However, this increased activation did not become more bilateral with task demand, thus providing no support for the hemispheric asymmetry reduction proposed by the HAROLD model (Cabeza, 2002).

When examining the relationship between brain activity and performance, distinct patterns emerged across difficulty levels. At lower difficulty, increased activity in subcortical regions was associated with better performance. In contrast, at higher difficulty, increased activation in cortical motor areas such as SMA, LPMd, and M1 was linked to poorer performance (Fig. 2.C). This pattern suggests a shift in functional relevance of subcortical versus cortical recruitment with increasing task demand. Such a shift is consistent with prior research on functional reorganisation in older adults (Michely et al., 2018; Tscherpel et al., 2020), but it contrasts with the posterior-to-anterior shift proposed by PASA (Davis et al., 2008). This pattern may reflect reduced reliance on automatised processing in older adults. Subcortical regions are typically associated with lower-effort, well-learned motor responses, while cortical regions remain engaged when actions are less familiar or still being learned (Penhune and Doyon, 2002). Therefore, the increased cortical recruitment observed here may alternatively reflect increased cognitive effort required to perform a complex task that has not yet become automatic, rather than inefficient processing. Previous work has demonstrated that older adults rely more on cortical than subcortical regions during motor-related tasks (Mouthon et al., 2018; Santos Monteiro et al., 2017). While this pattern was observed during the planning phase of bimanual movement and remained stable following training (Santos Monteiro et al., 2017), the present results extend this work by showing that the relationship between brain activity and performance differs across regions as task complexity increases. Building on this, we propose a novel functional reorganisation hypothesis: the Subcortical to Cortical Shift (SCOS). This hypothesis builds on CRUNCH, in that both describe increased recruitment of neural resources with rising task demands. However, SCOS specifically characterises a shift in the location of recruitment from subcortical to cortical motor areas across difficulty. Importantly, this interpretation is based on differences in performance-related activity rather than direct evidence of a shift in overall activation between regions.

The current study showed that when task demands were high, overactivation of core motor areas was linked to poorer performance, supporting previous studies that interpret such patterns as signs of inefficient rather than compensatory recruitment (Knights et al., 2021; Morcom and Henson, 2020). Adding further support to this, our results show participants who showed greater increases in activity in M1, SMA, PMd, and DLPFC across difficulty levels also exhibited larger declines in performance (Fig. 2.D). This directly contradicts the compensatory prediction from hypothesis 3 and instead strengthens the interpretation that increased activity in these regions reflects neural inefficiency rather than successful adaptation. Alternatively, rather than reflecting neural inefficiency, the increased activation may indicate that participants were engaging more cognitive effort to manage complex task rules that had not yet become familiar or automatic. However, this effort did not prevent the observed decline in performance.

Some limitations should be acknowledged. First, because of the absence of a younger control group (even though pre-planned) it remains possible that younger adults may show similar effects. Nonetheless to date only a handful of studies have directly examined the relationship between performance and modulation in older adults during cognitive-motor tasks (Santos Monteiro et al., 2020), and none were designed with older adults serving as their own controls. Here, in examining how neural activity changes with increasing difficulty within the same individuals, the present study provides a direct test of (or a lack of) compensatory capacity. Second, it is possible that the highest task difficulty represented the limit of the participants’ compensatory capacity. According to CRUNCH, compensatory mechanisms become ineffective once task demands exceed available neural resources. Thus, the performance decline observed at the highest difficulty may reflect participants reaching this compensatory ceiling, rather than indicating that compensatory processes were absent. Future studies should better characterise this potential ceiling effect, by including additional intermediate levels of difficulty, allowing a more detailed assessment of how neural recruitment changes across task demands. Finally, the assumption of a linear change in activity and performance across conditions had to be made, due to FSL’s constraint to linear slope modelling. This may have oversimplified the nature of the neural modulation, where other software, modelling nonlinear dynamics, may offer a more accurate representation of how older adults adjust to increasing demand.

## 5. CONCLUSION

This study provides evidence that older adults show increased brain activity in response to rising task complexity during cognitive-motor performance. However, this increased activity, particularly in cortical motor areas, was not associated with improved performance. Instead, subcortical activity supported performance at low difficulty, while greater cortical motor recruitment at higher difficulty was linked to poorer performance. These findings challenge compensation-based interpretations of PASA and HAROLD within the context of increasing task demands, suggesting that increased recruitment does not necessarily support performance under high cognitive-motor load. Instead, our results are more consistent with CRUNCH, where neural recruitment increases with task demand but becomes ineffective at higher levels of difficulty. Here we propose a shift in the location of the recruitment: from subcortical to cortical motor areas, as demand increases. This shift may characterise task demand-related neural reorganisation in older adults during complex motor decision-making. Furthermore, increased activity across difficulty levels in motor and prefrontal regions was linked to worse performance, opposing compensatory assumptions.

Overall, this study underscores the importance of examining how neural activity adapts dynamically to increases in task demands. It provides new insight into how cognitive-motor processing reorganises with increasing complexity in older adults and highlights the value of modulation-based approaches for understanding compensatory capacity.

## 6. FUNDING

This work was supported by the Economic and Social Research Council from the Scottish Graduate School of Social Science to MK (grant ref: ES/P000681/1) and by a Wellcome Career Development Award to CSB (ref: 320009/Z/24/Z).

## 7. ACKNOWLEDGEMENTS

The authors would like to thank the radiographer Frances Crabbe for her assistance with data collection, the University of the Third Age for support with participant recruitment, and all study participants for generously contributing their time.

For the purpose of open access, the author(s) has applied a Creative Commons Attribution (CC BY) licence to any Author Accepted Manuscript version arising from this submission.

## 8. AUTHORS CONTRIBUTION

M.K contributed to the conceptualisation of the study, methodology, project administration, the investigation and formal analysis, and writing the original draft. S.R contributed to the investigation and formal analysis. M.H contributed to the conceptualisation of the study, supervision, reviewing and editing the original draft. C.SB contributed to the conceptualisation of the study, methodology, project administration, resources, supervision, reviewing and editing the original draft.

## 9. COMPETING INTEREST

The authors have no competing interests.

## Notes

### Competing Interest Statement

The authors have declared no competing interest.

