## Supplementary Materials for "When More Is Not Better: Increased Motor Cortex Recruitment In Older Adults Is Associated With Performance Decline During High-Demand Cognitive-Motor Tasks"

### S1. ACTION SELECTION DIFFICULTY PILOTING

Three increasing AS difficulties were created derived from the original AS task (O'Shea 2007). From the original AS rule, extra shapes or colours were added to complicate the rule (Fig. S1.A). The four difficulties (first one being the original task) were then piloted on six older adults (mean age = 65.1, sd = 8.4). Three of these were chosen. A repeated-measures ANOVA revealed a significant difference between difficulties ($F(1.85,9.25) = 5.28$), $p = 0.03$, Fig. S1.B). Bonferroni post-hoc tests revealed RT cost was significantly higher in difficulty 4 vs 1 (paired t-test, $t(5) = -13$, pcorr < 0.001, Fig. S1.B). No difference was found between any other difficulty pair (see Fig. S1 legend for exact results). Additionally, when looking at the data (Fig. S1.B), it was clear that difficulty 3 was not difficult enough compared to the others. Hence, difficulty 1, 2 and 4 were kept for the study (renamed 1, 2 and 3).

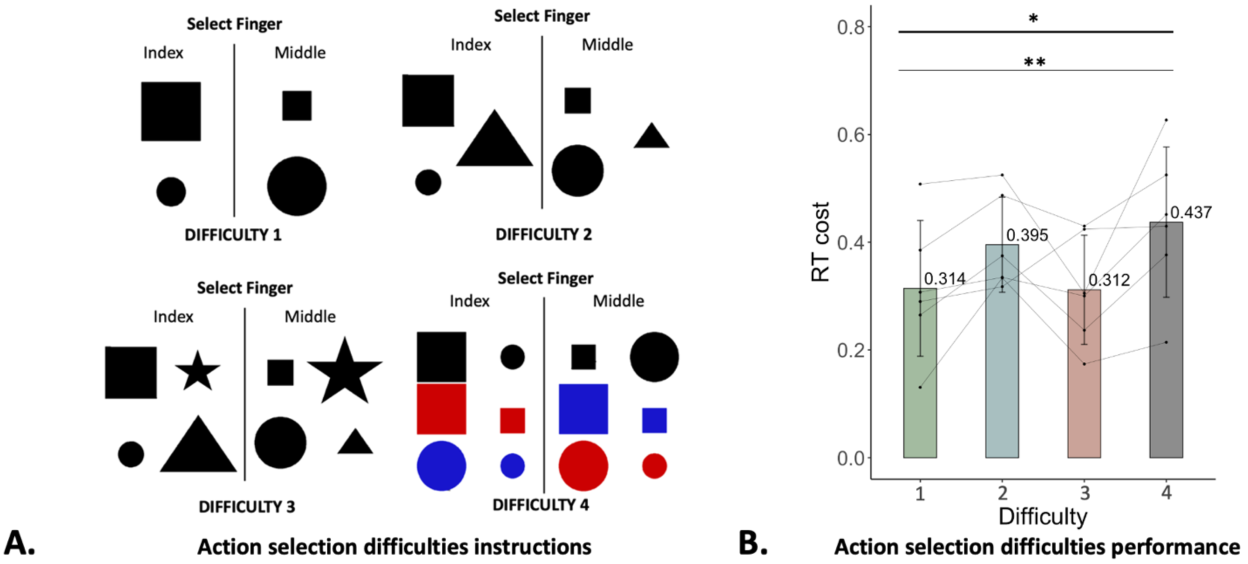


**Figure S1: AS difficulty piloting**

A. AS difficulties piloted. Compared to the first difficulty, a shape was added for difficulty 2 and two shapes for difficulty 3. For difficulty 4, colours (red and blue) were added.

B. AS performance (RT cost, sec) for each difficulty, N = 6.

Bonferroni post-hoc paired t-tests: 1 vs 2 $t(5) = -2.75$, pcorr = 0.12; 1 vs 3 $t(5) = 0.05$, pcorr = 1; 2 vs 3 $t(5) = -1.78$, pcorr = 1; 2 vs 4 $t(5) = -1.12$, pcorr = 0.94; 3 vs 4 $t(5) = -2.77$, pcorr = 0.12

### S2. ADDITIONAL ANALYSES

Exploratory analyses were carried out for each hypothesis as follows:

- Hypothesis 1, predicting that activity in SMA, M1, PMd, and DLPFC will increase with difficulty, was tested at a regional level.
- Hypothesis 2, stating that high activity in SMA, M1 and PMd will be linked to poorer performance in higher difficulty levels, and high activity in DLPFC with better performance; was tested at a whole-brain level using IES as a performance measure, and at a regional level with both RT cost and IES as performance measures.
- Hypothesis 3, predicting that greater activity increases in M1, LPMd, SMA and DLPFC across difficulty levels will be linked to lower performance decline across difficulty level, was tested at a whole-brain level using IES as a performance measure, and at a regional level using both RT cost and IES as performance measures.

#### S2.1 Methods

##### *S2.1.1 Region of interest definition*

ROIs were identified in MNI space, from a whole-brain BOLD mean analysis of the three difficulties. The left M1 (LM1) was identified as the peak activity around the hand knob representation area, during EO > VO contrasts in all three difficulties (x = -36, y = -24, z = 54, Fig. S2). Left PMd (LPMd) and SMA were located by finding the peak activity around the LPMd and SMA (Brodmann area 6) regions during the AS > EO in all three difficulties (LPMd: x = -26, y = -8, z = 56, SMA: x = 0, y = 16, z = 50, Fig. S2). Left DLPFC (LDLPFC) was located by finding the peak activity around the left middle frontal gyrus (Brodmann area 45) during the AS > EO in difficulty 3. Finally, the left auditory cortex was used as a control area to compare findings to. It was located from the Juelich atlas (primary auditory cortex (x = -40, y = -30, z = 11, Fig. S2). Each ROI location was then expanded to 1cm diameter spheres using FSLmaths and registered to each participant's native space.

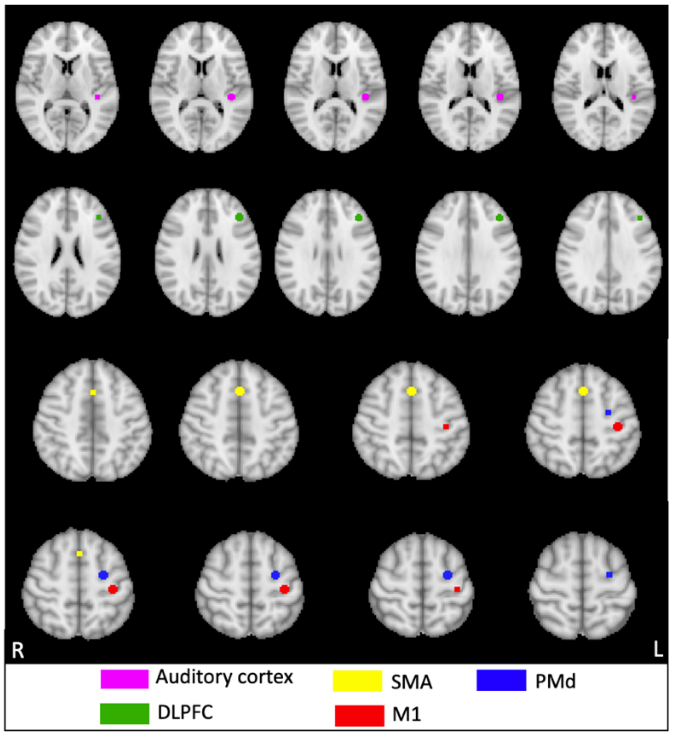


**Figure S2: Group ROI**

ROI were localised from whole-brain BOLD mean analysis of the three difficulties. Expanded to 1cm diameter spheres.

*S2.1.2 Regional activity analysis*

The percentage BOLD signal change was extracted during AS > EO contrast for each region (LPMd, SMA, LM1, LDPFC, left auditory cortex as a control area), using Featquery (Woolrich et al., 2004). This was done for each participant and difficulty separately.

To investigate main effects of difficulty (hypothesis 1), BH corrected one-way repeated-measures ANOVAs were conducted on SMA, LPMd and LDLPFC activity between difficulties. LM1 and left auditory cortex activity were not normally distributed (Shapiro-Wilk, LM1 $p = 0.03$; left auditory cortex $p < 0.0001$), hence BH corrected Friedman tests were conducted between difficulties. Bonferroni corrected post-hoc exploratory paired t-tests were conducted on significant main effects, between each difficulty.

To test hypothesis 2, the link between ROI activity and AS performance (IES and RT cost) was investigated. This was done using PALM (Winkler et al., 2014), conducting Pearson correlation for each difficulty separately, between AS performance (RT cost and IES) and each predicted region, as well as the control area (controlling for age). Both positive and negative correlations were tested using 1 and -1 contrasts.

Finally, to test hypothesis 3, the relationship between activity modulation and AS performance adaptation was investigated. For this, the linear slope of AS performance (IES and RT cost) over the three difficulties was calculated. This was also done for each region activity and the control area. PALM (Winkler et al., 2014) was then used to conduct Pearson correlation between AS performance slope and each region activity slope, as well as the control area (controlling for age). Both positive and negative correlations were tested using 1 and -1 contrasts.

A significance threshold of $p < 0.05$ was used.

*S2.1.3 PALM analysis statistics*

When running PALM for correlations with AS performance (RT cost and IES), threshold-free cluster enhancement and 10,000 permutations were used to determine positive and negative correlations. BH correction was then applied to the results across the regions and difficulties tested. A significance threshold of $p < 0.05$ was used.

**S2.2 Results**

*S2.2.1 AS performance (IES) declines with increasing difficulty*

As IES was not normally distributed across all difficulties (Shapiro-Wilk test: $p < 0.0001$) a Friedman test was used to test for the main effect of difficulty, followed by Bonferroni corrected post-hoc two-tailed Wilcoxon signed-rank tests between each difficulty.

A difference in IES between difficulties was tested for, as we expected to see a decrease in performance with increasing difficulty. This was confirmed, with a main effect of difficulty found (Friedman test $X^2 = 28.3$, $p < 0.001$, Fig. S3). When doing Bonferroni post-hoc tests, IES was significantly higher in difficulty 3 vs 1 (Wilcoxon paired signed-rank test, Z = - 4.76, pcorr < 0.0001, Fig. S3), as well as in difficulty 3 vs 2 (Wilcoxon paired signed-rank test, Z = -3.94, pcorr < 0.001, Fig. S3). No difference was found between difficulty 1 and 2 (Wilcoxon paired signed-rank test, Z = -0.2, pcorr = 1).

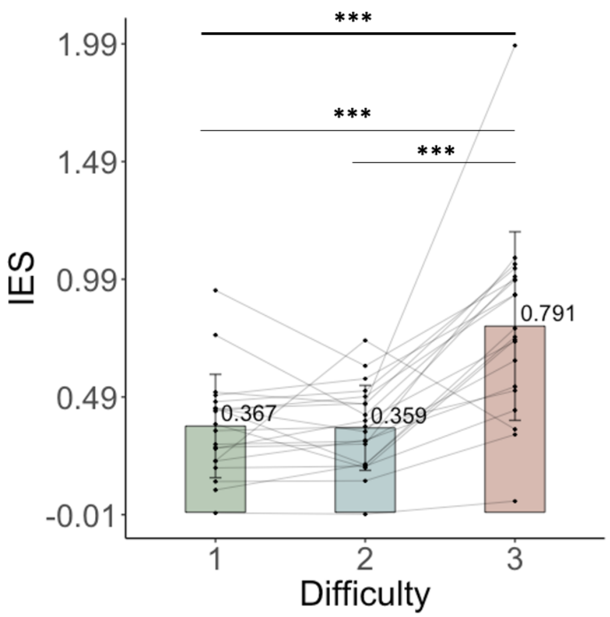


**Figure S3: AS performance between difficulties**

A.IES between difficulties. A high IES indicates a worse performance. Main effect of difficulty (Friedman test $X^2 = 28.3$, $p < 0.001$, N = 20).

Boxes represent means, error bars are standard deviations and dots are individual data points. Full thick lines illustrate significant main effects of difficulty, full thin lines represent significant post-hoc comparisons, dashed lines illustrate significant main effects that did not survive BH correction. Significant p-value of ≤ 0.001 represented by ***.

*S2.2.2 Sensorimotor and frontal networks activated during the AS task*

The activity present during AS was assessed for each difficulty, a lot of overlap was found between the difficulties. That is, for all three difficulties, positive activity was found in the SMA, bilateral superior and middle frontal gyrus (DLPFC), PMd, somatosensory cortex, frontal pole, insula, as well as LM1, paracingulate and supramarginal gyrus (all $p < 0.05$, corrected, Fig. S4.B.1). The precentral gyrus was found to be activated bilaterally in all difficulties but became spatially wider in the left hemisphere in difficulty 3. In addition, positive activity was found bilaterally in the superior parietal lobule and the thalamus in difficulty 1 and 3, as opposed to in the left hemisphere only for difficulty 2. Lastly, positive activations in the precuneus cortex were found in difficulty 3 only.

Opposed to that, negative activity was found in the left occipital cortex and subcallosal body in all three difficulties (all $p < 0.05$, corrected, Fig. S4.B.2). The cingulate gyrus and the precuneus cortex were found to be negatively activated during difficulty 1 and 2.

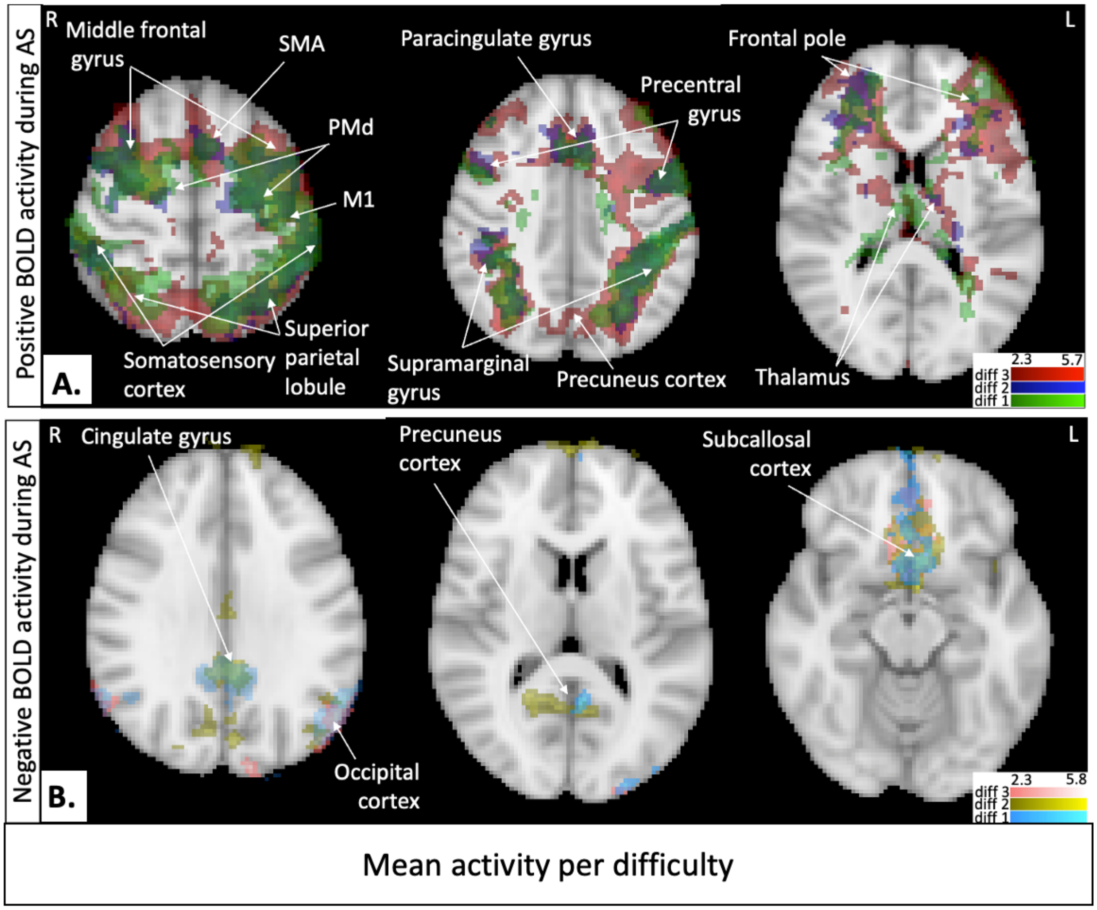


**<u>Figure S4:</u> Mean activity during AS task**
Analysis of whole-brain activity mean during AS > EO (all $p < 0.05$ corrected, N = 21 for difficulty 1 & 2, N = 20 for difficulty 3). A. Illustrates the positive BOLD activity (difficulty 1 in green, 2 in blue and 3 in red), whereas B. Illustrates the negative BOLD activity (difficulty 1 in cyan, 2 in yellow, 3 in pink).

*<u>S2.2.3 Increases in LPMd and LDLPFC activity with difficulty</u>*

To test hypothesis 1 using a ROI approach, ROI activity was investigated between difficulties and compared to a control area (left auditory cortex). A BH corrected repeated-measures ANOVA revealed a main effect of difficulty in LPMd ($F(2,38) = 5.42$, $p = 0.009$, pcorr = 0.02, Fig. S5.A.) and LDLPFC ($F(2,38) = 28.6$, $p = 0.001$, pcorr = 0.005, Fig. S5.B.), with both areas having increased activity at higher difficulty. A main effect of difficulty was found in SMA activity but did not survive BH correction ($F(2,38) = 3.72$, $p = 0.034$, pcorr = 0.057, Fig. S5.C.). BH corrected Friedman tests revealed no main effect of difficulty on LM1 activity ($X^2 = 2.1$, $p = 0.35$, pcorr = 0.44 Fig. S5.D), nor the control auditory cortex activity ($X^2 = 1.3$, $p = 0.52$, pcorr = 0.52, Fig. S5.E).

When doing Bonferroni corrected post-hoc paired t-tests on the significant main effects, LPMd activity was significantly higher in difficulty 3 vs 2 (t(19) = -3.1, pcorr = 0.018, Fig. S5.C.), but not between difficulty 1 and 2 (t(19) = 0.84, pcorr = 1), nor between difficulty 1 and 3 (t(19) = -2.27, pcorr = 0.11). Lastly, LDLPFC was found to be significantly higher in difficulty 3 vs 1 (t(19) = -3.54, pcorr = 0.007, Fig. S5.D), and in difficulty 3 vs 2 (t(19) = -3.35, pcorr = 0.01, Fig. S5.D) but no difference between difficulty 1 and 2 (t(19) = -0.02, pcorr = 1).

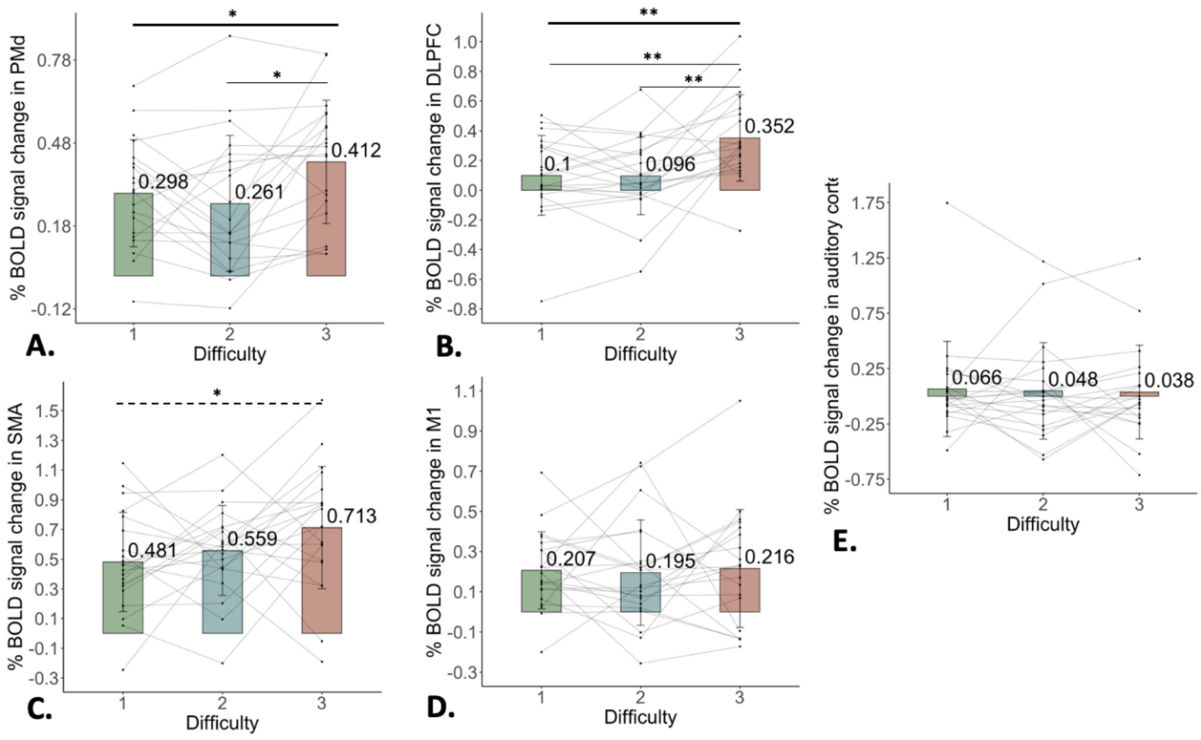


<u>**Figure S5: ROI activity per difficulty**</u>

Percentage BOLD signal change for each region (LPMd in A., LDLPFC in B., SMA in C., LM1 in D., and the control left auditory cortex in E.). Difficulties are represented in different colours (green for 1, blue for 2, red for 3). A BH corrected repeated-measures ANOVA showed a main effect of difficulty in LPMd (F(2,38) = 5.42, pcorr = 0.02) and LDLPFC (F(2,38) = 28.6, pcorr = 0.005). Individual bar charts represent means, error bars are standard deviations and dots are individual data points, N = 20. Full thick lines illustrate significant main effects of difficulty, full thin lines represent significant post-hoc comparisons, dashed lines illustrate significant main effects that did not survive BH correction. Significant p-value of ≤ 0.05 represented by *, ≤ 0.01 represented by **.

*S2.2.4 Motor planning areas linked to better AS performance at low difficulty and poorer performance at high difficulty*

When investigating whole-brain activity correlation with AS performance measured by IES for hypothesis 2, the bilateral frontal pole, caudate nucleus and right putamen (subparts of the basal ganglia), as well as the middle temporal gyrus were found to be correlated with better performance in difficulty 1 (all $p < 0.05$, corrected, Fig. S6.A.1). In difficulty 2, better performance was correlated with activity in the left occipital cortex (Fig. S6.A.1). No brain areas were found to be correlated with better performance in difficulty 3.

Opposed to that, poorer performance in the AS task (high IES) was correlated with activity in the left frontal pole and occipital cortex in difficulty 1 (all $p < 0.05$ corrected, Fig. S6.A.2). In difficulty 2, activity in the left frontal pole was associated with poorer performance (all $p < 0.05$ corrected, Fig. S6.A.2). Lastly, poorer performance was associated with activity in the SMA, LPMd, inferior parietal lobule, the right superior frontal gyrus and the bilateral secondary somatosensory cortex in difficulty 3 (all $p < 0.05$ corrected, Fig. S6.A.2).

*S2.2.5 No relationship between ROI activity and AS performance*

Hypothesis 2 was then tested at ROI level using a ROI-based approach. As there was no sign of increased recruitment of the right hemisphere at higher difficulty in the whole-brain analysis (Fig. S3.B main results), the ROI analysis only focused on the left hemisphere. After correcting for multiple comparisons across regions and difficulties, no significant relationship was found between SMA, LDLPFC, LPMd or LM1 and IES (Fig. S6.B.1) or RT cost (Fig. S6.B.2) in any of the difficulties. Similarly, no correlation was present with the control auditory cortex activity. The correlations coefficients and p-values are reported in Fig. S6.B.

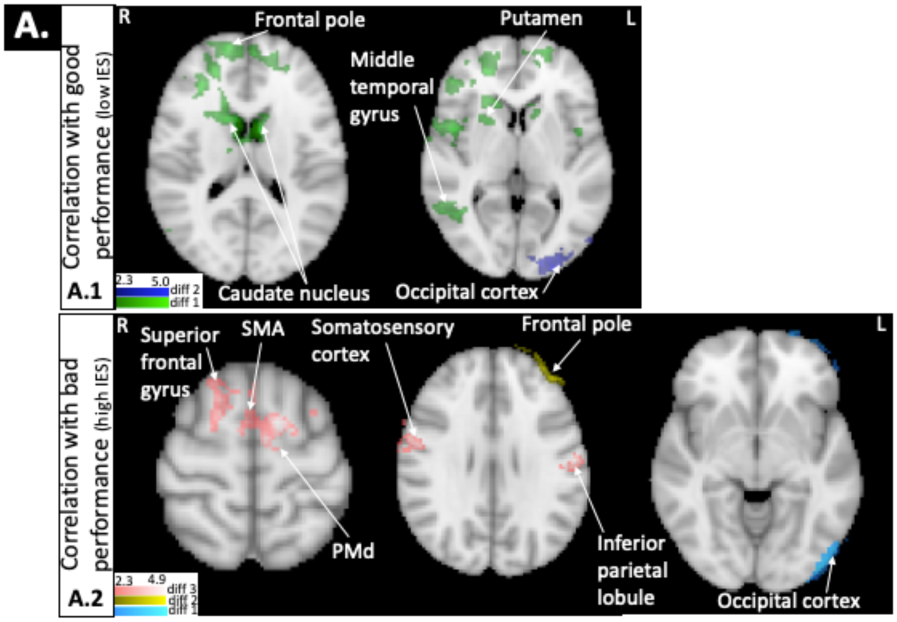


**B.**

**Correlation with IES**

**B.1**

| | DIFFICULTY 1 | | | DIFFICULTY 2 | | | DIFFICULTY 3 | | |
|---|---|---|---|---|---|---|---|---|---|
| **ROI** | **Pearson r** | **p-value** | **Corrected p-value** | **Pearson r** | **p-value** | **Corrected p-value** | **Pearson r** | **p-value** | **Corrected p-value** |
| **SMA** | 0.067 | 0.381 | 0.44 | 0.208 | 0.173 | 0.368 | 0.356 | ***0.046*** | 0.368 |
| **DLPFC** | -0.142 | 0.237 | 0.368 | 0.198 | 0.199 | 0.368 | -0.271 | 0.112 | 0.368 |
| **PMd** | -0.088 | 0.35 | 0.438 | 0.213 | 0.175 | 0.368 | 0.342 | 0.065 | 0.368 |
| **M1** | -0.037 | 0.44 | 0.471 | 0.012 | 0.477 | 0.477 | 0.314 | 0.094 | 0.368 |
| **Auditory cortex** | -0.138 | 0.27 | 0.368 | -0.153 | 0.268 | 0.368 | 0.202 | 0.189 | 0.368 |

**Correlation with RT cost**

**B.2**

| | DIFFICULTY 1 | | | DIFFICULTY 2 | | | DIFFICULTY 3 | | |
|---|---|---|---|---|---|---|---|---|---|
| **ROI** | **Pearson r** | **p-value** | **Corrected p-value** | **Pearson r** | **p-value** | **Corrected p-value** | **Pearson r** | **p-value** | **Corrected p-value** |
| **SMA** | 0.099 | 0.332 | 0.424 | 0.084 | 0.359 | 0.424 | 0.439 | ***0.023*** | 0.173 |
| **DLPFC** | -0.129 | 0.294 | 0.424 | 0.163 | 0.24 | 0.424 | -0.218 | 0.173 | 0.371 |
| **PMd** | -0.064 | 0.394 | 0.424 | 0.391 | ***0.041*** | 0.205 | 0.307 | 0.091 | 0.341 |
| **M1** | 0.227 | 0.165 | 0.371 | 0.246 | 0.133 | 0.371 | 0.512 | ***0.007*** | 0.105 |
| **Auditory cortex** | 0.046 | 0.447 | 0.447 | -0.062 | 0.396 | 0.424 | 0.106 | 0.328 | 0.424 |

**Figure S6: Whole-brain activity relationship with AS performance**

Correlation analysis between whole-brain activity (AS > EO) and AS performance (IES in A., RT cost in B.) (all $p < 0.05$ corrected, N = 21 for difficulty 1 & 2, N = 20 for difficulty 3). BOLD activity correlated with better performance (low IES) is illustrated in A.1 (green gradient for difficulty 1, blue for difficulty 2). BOLD activity correlated with poorer performance (high IES) is illustrated in A.2 (cyan gradient for difficulty 1, yellow for difficulty 2, pink for difficulty 3).
BOLD activity correlated with better performance (low RT cost) is illustrated in B.1 (green gradient for difficulty 1). BOLD activity correlated with poorer performance (high RT cost) is illustrated in B.2 (cyan gradient for difficulty 1, yellow for difficulty 2, pink for difficulty 3).
Relationship between AS performance and ROI BOLD activity (SMA, LDLPFC, LPMd, LM1, and the control left auditory cortex). AS performance measured with IES in A. and with RT cost in B. Analysis done

using PALM Pearson correlation, controlling for age. Multiple comparison correction was done using Benjamin-Hochberg method across regions and difficulties. N = 21 for difficulty 1 and 2, N = 20 for difficulty 3.

Overall, areas involved in motor planning (basal ganglia subparts) were linked to better performance in low difficulties compared to other areas with similar functions (SMA, LPMd, LM1) being linked to poorer performance in high difficulties, partly in line with the first part of the hypothesis but not replicated in the ROI analysis.

#### *S2.2.6 M1 activity modulation with difficulty related to better AS performance adaptation*

When testing for hypothesis 3, at a whole-brain level, no correlation (positive or negative) was found between activity slope and IES slope. This suggests no relationship between changes in activity over difficulties and performance adaptation (measured with IES).

When investigated at a regional level, the relationship between changes in ROI BOLD activity with increasing difficulty and changes in AS performance (IES slope and RT cost) were investigated. Change was quantified by calculating the linear slope of activity and IES across difficulty levels. After correcting for multiple comparisons, a negative relationship between the LM1 activity slope and the IES slope was found ($r = -0.561$, $p_{corr} = 0.02$, Fig. S7.A). This suggests that, with increasing difficulty, the greater increase in LM1 activity was linked to less decline in AS performance (measured by IES). No significant relationship was found for the other region or the control auditory cortex (Fig. S7.A.1 for p-values).

When using RT cost as a AS performance measure, no significant relationship was found between any of the regions (Fig. S7.B for p-values).

There were no significant effects for the control auditory cortex activity slope and AS performance slope (IES or RT cost) (Fig. S7.B for p-values).

Overall, the LM1 activity increases seemed to be beneficial for the adaptation of the AS performance at high difficulty (measured by IES only), implying a potential compensatory role. This finding contrasts with the whole-brain analysis, which showed that greater motor activation was generally associated with poorer performance. Importantly, the LM1 ROI used in the regional analysis was located a few millimetres away from the peak motor region identified in the whole-brain correlation map, which may account for this discrepancy.

**A.** **Correlation with IES slope**

**A.1**

| ROI activity slope | Pearson r | p-value | Corrected p-value |
|---|---|---|---|
| **SMA** | -0.218 | 0.16 | 0.245 |
| **M1** | -0.561 | ***0.004*** | ***0.02 **** |
| **PMd** | -0.197 | 0.196 | 0.245 |
| **DLPFC** | 0.101 | 0.313 | 0.313 |
| **Auditory cortex** | -0.231 | 0.153 | 0.245 |

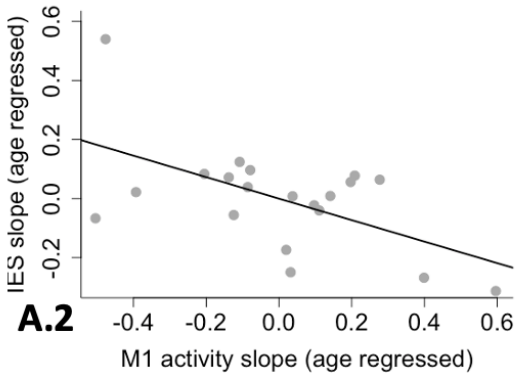


**B.** **Correlation with RT cost slope**

| ROI activity slope | Pearson r | p-value | Corrected p-value |
|---|---|---|---|
| **SMA** | -0.191 | 0.198 | 0.335 |
| **M1** | -0.213 | 0.179 | 0.335 |
| **PMd** | 0.085 | 0.358 | 0.358 |
| **DLPFC** | 0.118 | 0.291 | 0.358 |
| **Auditory cortex** | -0.193 | 0.201 | 0.335 |

**Figure S7: Relationship between activity modulation and performance adaptation**

Relationship between ROI activity slope and performance slope over the three difficulties (IES in A., RT cost in B.) for each ROI (SMA, LM1, LPMd, LDLPFC and the control left auditory cortex). Analysis done using PALM Pearson correlation, controlling for age. Multiple comparison correction was done using Benjamin-Hochberg method across region, N = 20. A negative correlation indicates that, with increasing difficulty, the greater increase in activity is linked to less decline in performance (lower increase in IES or RT cost). The significant negative correlation between IES slope and M1 activity slope is represented in A.2
